# A Case-Based Explainable Graph Neural Network Framework for Mechanistic Drug Repositioning

**DOI:** 10.1101/2025.04.28.651120

**Authors:** Adriana Carolina Gonzalez-Cavazos, Roger Tu, Meghamala Sinha, Andrew I Su

## Abstract

Drug repositioning offers a cost-effective alternative to traditional drug development by identifying new uses for existing drugs. Recent advances leverage Graph Neural Networks (GNN) to model complex biological data, showing promise in predicting novel drug-disease associations. However, these frameworks often lack explainability, a critical factor for validating predictions and understanding drug mechanisms. Here, we introduce Drug-Based Reasoning Explainer (DBR-X), an explainable GNN model that combines a link prediction module and a path-identification module to generate interpretable and faithful explanations. When benchmarked against other GNN link prediction frameworks, DBR-X achieves superior performance in identifying known drug-disease associations, demonstrating higher accuracy across all evaluation metrics. The quality of DBR-X biological explanations was assessed through multiple approaches: comparison with manually-curated drug mechanisms, evaluation of explanation faithfulness through deletion and insertion studies, and measurement of stability under graph perturbations. Together, our model not only advances the state-of-the-art in drug repositioning predictions but also provides multi-hop explanations that can accelerate the translation of computational predictions into clinical applications.

## 1 Introduction

Traditional drug discovery and development is a time-consuming and resource-intensive process [1]. To address these challenges, drug repositioning has emerged as a strategic approach to reduce development costs, lower failure rates, and expedite time to market [2]. Drug repositioning strategy is based on the premise that existing drugs may exhibit effects beyond their original targets. Despite its promise, most successful drug repositioning instances are often based on clinical observations. Hypothesis-driven identification of new applications for drug candidates remains problematic due to the complexity and limited understanding of the underlying mechanisms, as well as the dispersed nature of relevant information in a growing sea of information.

A recent area in computational drug repositioning leverages knowledge graphs (KGs), a data structure that models relationships between various biomedical entities like drugs, genes, diseases, etc., in the form of triplets [3–7]. By representing biological entities as nodes and their associations as edges, KGs allow for the integration and analysis of complex biological data — facilitating the discovery of hidden connections and predicting potential new uses for existing drugs. While Graph Neural Networks (GNNs) have emerged as powerful tools for analyzing these multi-relational structures and predicting novel therapeutic applications, their lack of interpretability limits their practical utility in drug discovery [8–12].

Recent advances in Case-Based Reasoning (CBR) have shown promise in addressing this interpretability challenge. CBR is an artificial intelligence framework that learns from past experiences to solve new problems through three main steps: retrieving similar cases to the given problem, reusing their solutions, and revising if the solutions are appropriate [13]. In knowledge graph applications, Das et al. pioneered a non-parametric CBR approach for KG completion, finding reasoning chains for queries by retrieving similar cases [14]. While innovative, this approach was limited to finding similarities in patterns between cases and queries using simple symbolic string matching. A follow-up work proposed CBR-SUBG, a semi-parametric model that leverages similarities in local subgraph structures to answer queries [15]. However, while the CBR-SUBG framework outperforms baseline link-prediction models on benchmark datasets, it lacks the ability to highlight the important connections that explain their predictions.

Several explainable AI techniques have been proposed to generate explanations for GNN predictions, mainly differing in the search method employed within the subgraph space. GNNExplainer pioneered this field by introducing parameterized masks for both edges and node features, maximizing mutual information between masked subgraphs and original predictions [16]. PGExplainer later adopted a deep neural network to parameterize the explanation generation process, enabling collective explanation of multiple instances [17]. However, these methods can produce disconnected subgraphs without structural constraints. While PaGE-Link attempted to solve this by focusing on path-based explanations using a heterogeneous path-enforcing mask, its pruning approach, which removes nodes based on their connectivity and eliminates nodes multiple hops away from the source or target, can lead to incomplete or inaccurate explanations [18].

In this work, we introduce Drug-Based Reasoning Explainer (DBR-X), a framework that provides path-based explanations for drug-disease predictions through two complementary modules: (i) a link prediction module that identifies potential drug-disease connections by recognizing similar subgraph patterns using a CBR approach, and (ii) a path identification module that employs a heterogeneous path-enforcing mask with node degree scoring to identify mechanistically relevant connections. Unlike other GNN explainers that identify important subgraphs without domain context, DBR-X leverages similar indication cases as templates and identifies important multi-hop paths with optimal connectivity patterns, avoiding both overly-connected hub nodes and sparsely connected nodes that may represent noise.

We trained DBR-X on a biomedical knowledge graph to predict known drug-disease candidates and provide path-based explanations. Our evaluation shows that DBR-X outperforms other GNN-link prediction frameworks as well as state-of-the-art GNN explainer baselines. The quality of top-ranked connections was assessed across multiple performance axes, including faithfulness (via deletion and insertion) and stability. Identified important connections were also validated against a external gold-standard drug mechanisms dataset. Lastly, through detailed case studies, we show how DBR-X can identify promising drug repositioning candidates for rare diseases and provide biologically plausible rationales that can guide expert evaluation and experimental validation.

## 2 Results

### 2.1 Case-Based Reasoning for drug discovery: retrieving and adapting similar drug cases

The DBR-X model comprises two integrated modules: (i) a link prediction module that identifies potential drug-disease associations by retrieving similar cases, and (ii) a path identification module that elucidates mechanistic insights by delineating biologically plausible pathways connecting drugs to diseases.

The link prediction module employs a CBR approach through a three-step workflow (Figure 1a). For a query such as (Pranlukast, *indication*, ?), DBR-X first **retrieves** *k* - nearest neighbor (*k*-NN) drug entities (e.g., astemizole, sulindac, ketotifen) using a pre-computed similarity matrix derived from drug connectivity patterns in a knowledge graph (see Methods). For each retrieved drug, multi-hop paths to associated diseases are collected. These paths are then **reused** by applying their meta-path structures to the query drug, generating a query subgraph that suggests potential mechanisms and disease candidates. Finally, DBR-X utilizes a graph neural network (GNN) to **revise** and compare the local structural features of the query and *k*-NN drug subgraphs.

**Fig. 1.**
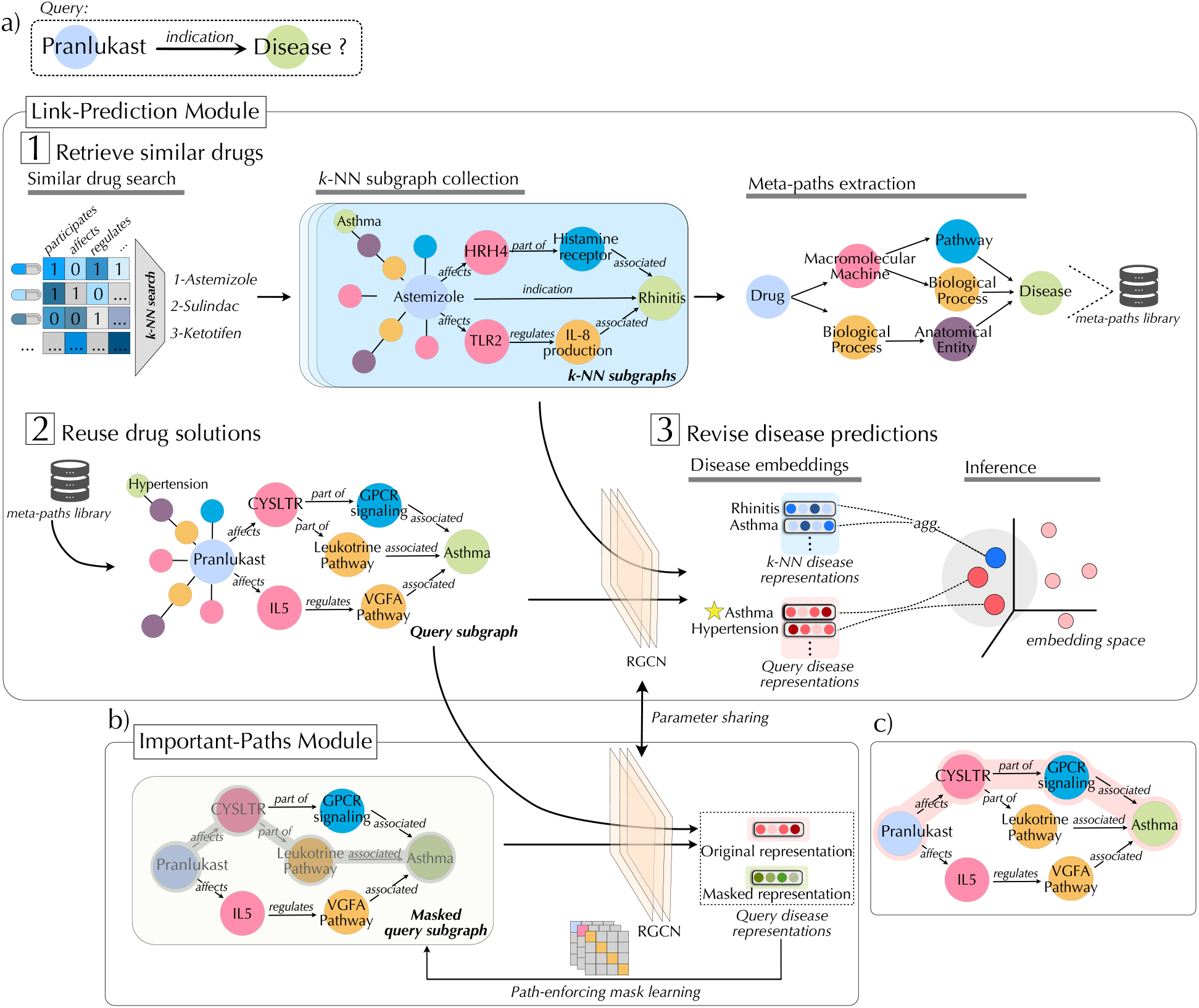
Drug repositioning predictions and important paths produced by Drug-based reasoning explainer (DBR-X) framework. A) Link-prediction module. (1) Given a query (Pranlukast, *indication*, ?), DBR-X first retrieves k-NN similar drug entities (astemizole, sulindac, ketotifen) and gathers their reasoning paths that lead to the corresponding disease they treat. (2) Next, reasoning paths are reused to address the query of interest. (3) Finally, disease answers gathered on query subgraph are revised. The answer node can be found by identifying the disease in the query subgraph that is most similar to the answer nodes in the subgraph of *k*-NN similar drug entities (marked with a star). Note that the corresponding answer node is analogously located within the reasoning patterns in each subgraph. B) Important-paths module. The module employs a heterogeneous path-enforcing mask with node degree scoring to identify a set of crucial paths that best explain the drug-disease relationship. The disease representations generated from the masked and original subgraphs are compared, allowing DBR-X to iteratively learn which edges are crucial for maintaining the original disease representation learned on link-prediction module. C) DBR-X identifies interpretable path explanations. Final explanatory output for the Pranlukast-asthma prediction, focusing on the most relevant (red) multi-hop path selected by DBR-X.

Grounded in the CBR principle that similar cases yield analogous outcomes, DBR-X assesses the GNN-generated representations of disease nodes in the query subgraph against those in the *k*-NN subgraphs. It then produces a ranked list of disease targets, with rankings reflecting the similarity between disease node representations in the query subgraph and those of known therapeutic targets in similar drug subgraphs. This approach enables DBR-X to prioritize the most promising disease targets for the query drug.

Visualization of the DBR-X embedding representation shows that the model effectively captures both global entity relationships and local structural similarities. In Figure 2a, the Uniform Manifold Approximation and Projection (UMAP) visualization of entity embeddings displays distinct clusters corresponding to biological entity types, indicating that DBR-X captures meaningful representations that reflect the hierarchical organization of the biomedical knowledge graph.

**Fig. 2.**
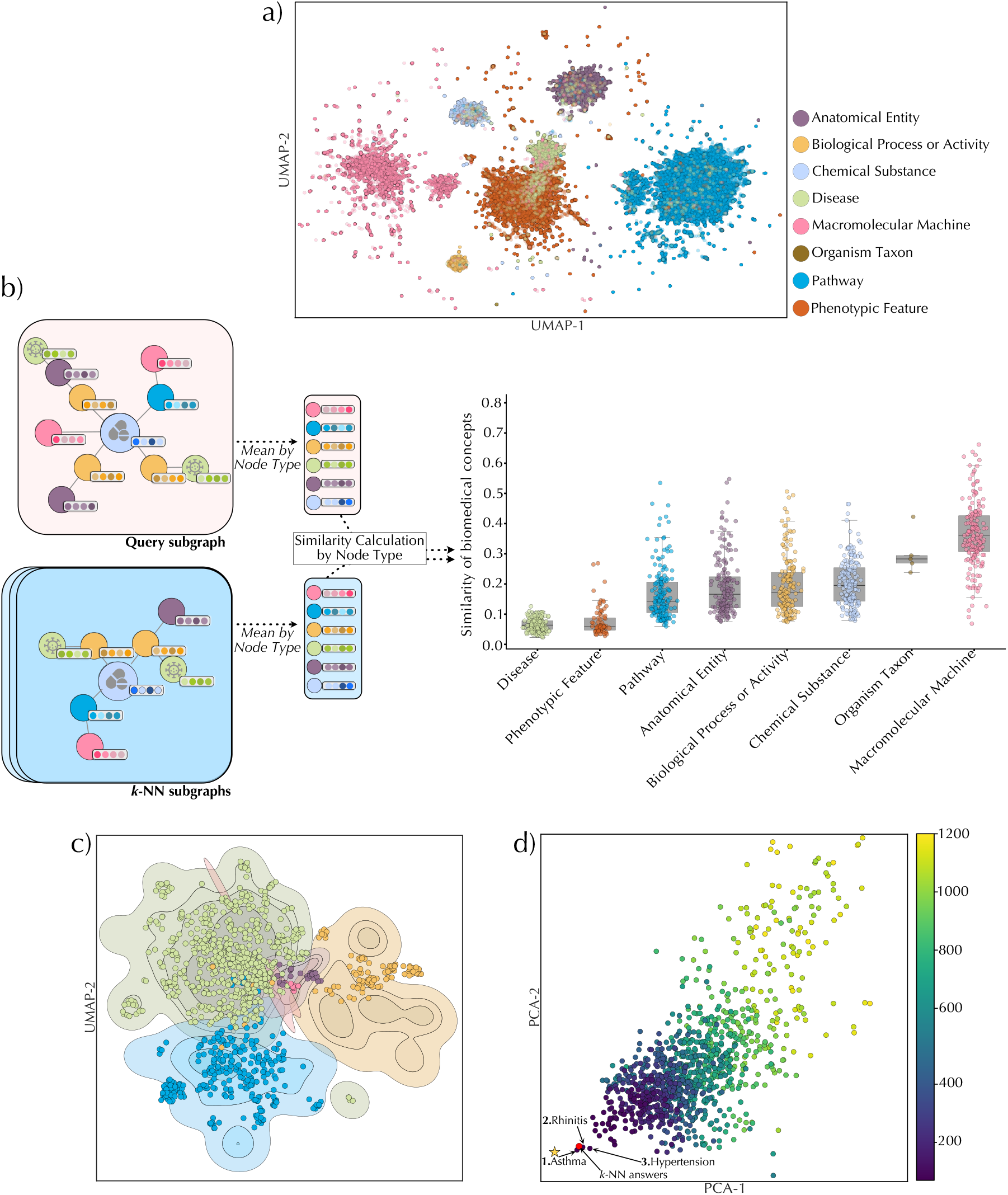
The embedding space of the DBR-X model. A) UMAP visualization of entity embeddings learned by DBR-X across the knowledge graph, with nodes colored by node type. B) DBR-X preserves structural similarities between query subgraph and retrieved subgraph drug cases. The left panel shows a representative query subgraph an6d its corresponding *k*-NN subgraphs retrieved by DBR-X. For each node type, we averaged all node representations in the query and compared this to the average of the same type in similar cases. The right panel presents the distribution of Euclidean distances comparing entity representations between query subgraphs and similar drug cases, grouped by biological entity type. Lower distance scores mean greater similarity between the query and its neighbors. C) UMAP projection of biomedical entities within Pranlukast query subgraph (dot) and corresponding similar entities (represented as contour plot). D) Disease prediction space for Pranlukast visualized using PCA. The mean representation of disease nodes from k-NN similar drugs is shown as a red dot. The color gradient represents the repositioning ranking score. The figure highlights the expected true disease (star) and the top predicted diseases.

We further assessed how DBR-X preserves subgraph structural similarities among similar drug cases (Figure 2b). To assess similarity, we calculated the Euclidean distances between averaged node representations for each biological entity type (e.g., drugs, genes, pathways) within the query subgraph and the corresponding averages in the k-NN subgraphs. The results revealed that DBR-X consistently maintains stable representations for higher-level biological entities (pathways and biological processes), successfully capturing the shared therapeutic mechanisms connecting similar drugs. Notably, we observed higher similarity variance specifically for macromolecular machine elements—an expected pattern that reflects established pharmacological principles, where drugs with similar therapeutic effects often work through different gene targets while ultimately influencing the same biological pathways. Figure 2c then illustrates this pattern using Pranlukast as a case study, where we visualize biological entities from Pranlukast’s subgraph (shown as dots) overlaid with contour plots representing the distribution of entities from similar cases. The clear alignment between Pranlukast’s entities and the contour regions demonstrates how DBR-X successfully captures shared biological mechanisms between similar cases.

To identify potential novel repositioning candidates, DBR-X generates node representations on both the drug query subgraph and the retrieved subgraphs of similar drugs. Potential disease targets are then ranked based on how closely their representations match those of known therapeutic targets from similar drug cases. Figure 2d illustrates this process using Pranlukast as a case study. The PCA projection shows the disease prediction space, where the mean representation of diseases from similar drug cases is marked with a red dot. The visualization reveals that asthma and rhinitis, Pranlukast’s known indication, clusters near this reference point [19].

Furthermore, the model identifies hypertension as top repositioning candidate for Pranlukast, as it also clusters close to the mean disease representation from similar cases. While no direct studies have examined Pranlukast’s effect on hypertension, it is reported that leukotriene antagonists can influence blood pressure regulation [20].

### 2.2 Performance of DBR-X predictions on known indications

Next, we assessed DBR-X’s ability to predict known FDA-approved indications using the Mechanistic Repositioning Network with Indications (MIND) KG dataset, comparing it against state-of-the-art knowledge graph completion models [21]. Specifically, we benchmark against GNN-based link prediction models, including Relational Graph Convolutional Network (R-GCN) [22] and composition-based multi-relational Graph Convolutional Network (CompGCN) [23]. The evaluation of GNN-based models was performed using ConvE [24] and DistMult [25] as scoring functions. We also compared DBR-X against the traditional non-parametric CBR method, which identifies drug-disease connections by matching existing path patterns between source and target entities without any learned parameters [14].

We evaluated DBR-X’s link prediction capabilities using two standard ranking metrics. The Hits@*K* metric, which measures how often the correct disease appears within the top *K* predictions, reflecting the model’s accuracy in ranking relevant disease associations. Mean reciprocal rank (MRR) calculates the average inverse position of correct disease predictions, providing an overall measure of prediction accuracy across all cases. As shown in Table 1, DBR-X demonstrates consistently superior performance across all evaluation metrics compared to GNN-baseline and non-parametric CBR methods, suggesting that DBR-X’s strategy of focusing on query-relevant subgraphs provides significant advantages over methods that process the complete knowledge graph.

**Table 1.** Link prediction performance

| Model | MRR | Hits@1 | Hits@3 | Hits@5 | Hits@10 |
| --- | --- | --- | --- | --- | --- |
| RGCN + DistMult | 0.1157 | 0.0180 | 0.1084 | 0.2108 | 0.3494 |
| RGCN + ConvE | 0.1197 | 0.0240 | 0.1144 | 0.1867 | 0.3614 |
| CompGCN + DistMult | 0.27735 | 0.1954 | 0.30651 | 0.36398 | 0.44828 |
| CompGCN + ConvE | 0.36112 | 0.25287 | 0.41762 | 0.47126 | 0.55556 |
| Non-parametric CBR | 0.03269 | 0.0158 | 0.0317 | 0.0396 | 0.0634 |
| DBR-X (ours) | **0.3770** | **0.2796** | **0.4329** | **0.4789** | **0.5708** |

### 2.3 Identifying biologically relevant paths through path-enforcing mask learning

The second module of DBR-X generates interpretable explanations by identifying relevant biological pathways that connect drugs to their predicted disease targets. This is accomplished through a heterogeneous path-enforcing mask learning approach (Figure 1b). For each drug-disease subgraph created by the link-prediction module, DBR-X identifies mechanistically meaningful paths using two learning objectives: a prediction-based loss that identifies influential edges for model prediction, and a path-based loss that ensures selected edges form biologically meaningful paths (see Methods).

The prediction-based loss utilizes a perturbation strategy to assess edge importance. It evaluates how masking specific edges from the query subgraph impacts the learned representations from the link-prediction module. For instance, if removing an edge significantly alters the embedding representation of the query disease node—thereby reducing its similarity to disease nodes from similar drug cases—that edge is deemed critical for maintaining predictive accuracy. These critical edges are assigned higher weights in the mask, reflecting their importance to the drug-disease association prediction. During mask learning, the initial masked representation of the query disease often differs substantially from the original representation. As training progresses, this difference diminishes, indicating that the mask successfully identifies edges essential for preserving the disease node’s original embedding. To ensure the mask identifies only the most essential edges, we include mask norm regularization, encouraging the mask to be discrete and sparse.

The path-based loss ensures that explanations are concise and biologically meaningful by imposing two key constraints. First, multi-hop explanations should avoid both highly connected hub nodes and extremely sparse nodes. To do this, DBR-X uses a scoring system (*D*_*s*_) that rates nodes based on their number of connections compared to an ideal value (*γ*) (see Methods). This helps pick pathways that go through nodes with a balanced number of links, making them more likely to reflect real biology instead of noise or overly general connections. Second, DBR-X restricts explanations to short paths, avoiding unnecessarily long or convoluted routes between drugs and diseases. DBR-X uses Dijkstra’s algorithm to find the clearest and shortest pathways in the weighted subgraph, ensuring the explanations are straightforward and practical.

### 2.4 Evaluation of DBR-X explanations against gold-standard drug mechanism

The ultimate goal of model explanation is to enhance transparency and support decision-making. For this, we evaluated our model’s explanatory power against drug mechanisms captured by DrugMechDB. To ensure a fair comparison, we modified baseline models to integrate CBR’s hypothesis-driven approach, resulting in three adapted versions: CBR+GNNExplainer, CBR+PGExplainer, and CBR+PaGE. This adaptation allowed the baseline models to focus on query-relevant subgraphs identified via case-based reasoning, rather than analyzing the entire graph structure. Additionally, since DBR-X and CBR+PaGE-Link models produce sets of paths, while CBR+GNNExplainer and CBR+PGExplainer highlight significant connections from a learned mask—we standardized our comparison using learned masks from all methods. In this framework, edges present in DrugMechDB ground truth paths were classified as positive, while all others were considered negative.

As shown in Figure 3a, DBR-X demonstrates superior ROC-AUC performance against our baselines when evaluated using the gold-standard DrugMechDB mechanisms [6]. For this evaluation, we considered edges from DrugMechDB paths as positive examples and all other edges as negative examples, then evaluated the mask weights (*M*) as prediction scores using the ROC-AUC metric. Here, a high ROC-AUC score reflects that edges in DrugMechDB ground truth mechanisms are precisely captured by the model mask. Here, a high ROC-AUC score reflects that edges in DrugMechDB ground truth mechanisms are precisely captured by the model mask. This substantial improvement is attributed to DBR-X’s node degree scoring, which ensures paths traverse through nodes with appropriate connectivity patterns (Figure 3b). By considering nodes with moderate degree values (*λ*=10), DBR-X effectively identifies mechanistically relevant paths while avoiding both overly-connected hub nodes that provide little specific information and extremely sparse nodes that may represent noise. This balanced approach to node selection, absent in baseline models, enables DBR-X to more accurately capture biologically meaningful mechanisms.

**Fig. 3.**
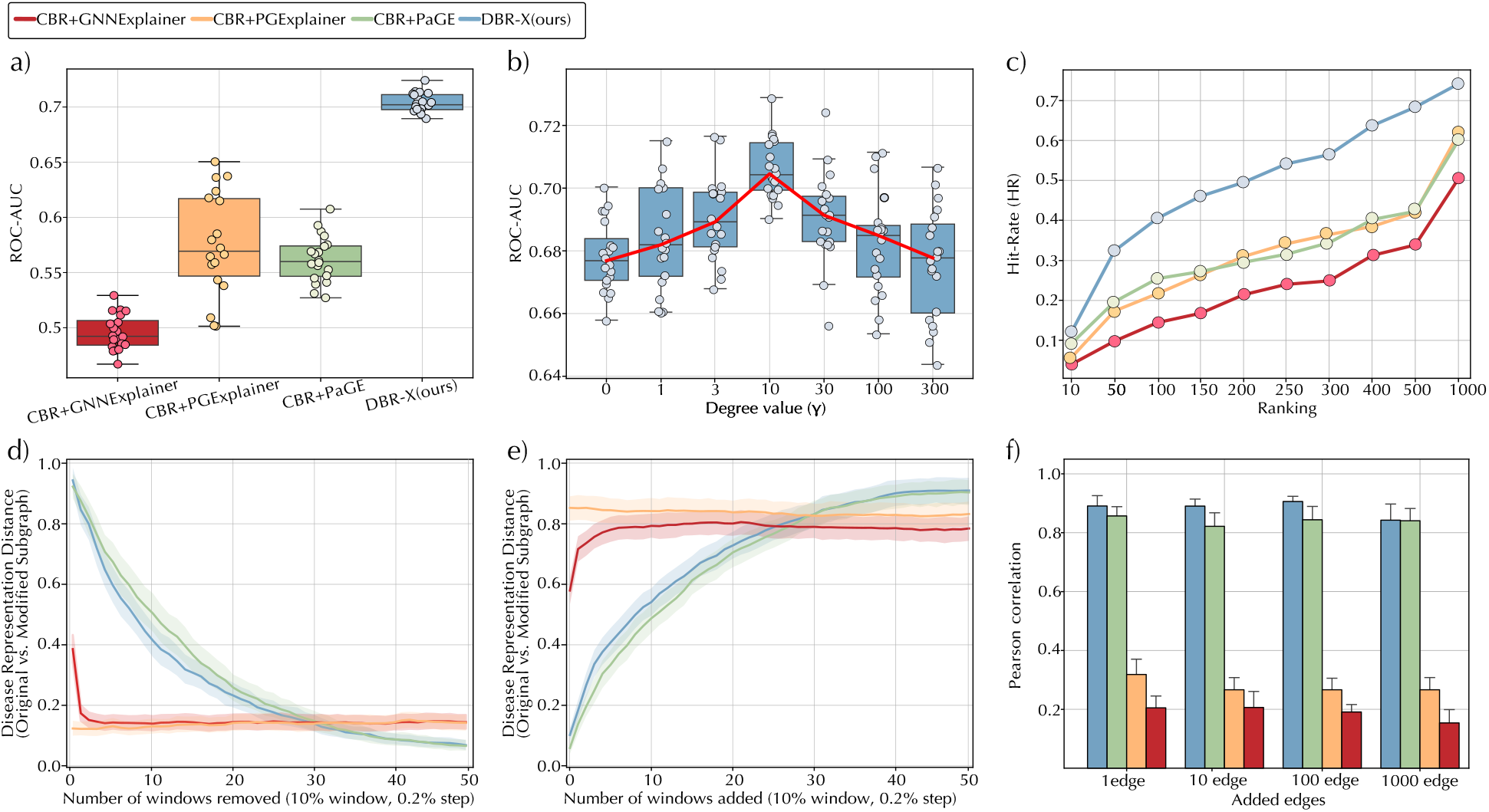
Evaluation of DBR-X important-paths module. A) Explanation accuracy: ROC-AUC scores comparing DBR-X and baseline models, where edges in DrugMechDB paths serve as ground truth. Mask weights are treated as edge prediction scores. B) Parameter sensitivity: ROC-AUC performance of DBR-X with varying optimal node degree values (*λ*), showing how this hyperparameter affects explanation quality. C) Edge Hit Rate: Evaluation at different ranking positions showing the proportion of ground truth edges captured at various thresholds. D) Deletion: Results show how much the model’s predictive performance degrades when edges, prioritized by their learned mask weights, are removed from the query subgraph. E) Insertion: Results show how much the model’s predictive performance maintains when edges, prioritized by their learned mask weights, are retained in the query subgraph. F) Stability: Robustness of explanations under random edge perturbations, measured by Pearson correlation between original and perturbed mask weights. Higher correlation values indicate more consistent explanations despite graph modifications. Note: All evaluations from a-b are conducted with N= 20. For d) and e) we use a sliding window, starting at the top 10% and shifting in 0.2% steps. Only the first 50 windows are displayed in results.

We further evaluated model performance through edge hit rate (HR@K) analysis, which measures the recovery rate of DrugMechDB’s ground truth edges at different ranking thresholds. As shown in Figure 3c, DBR-X substantially outperformed all baseline approaches, achieving hit rates nearly twice those of existing methods across all K values. While baseline models struggled to identify relevant edges even at higher K values, stabilizing at HR values between 0.35-0.4, DBR-X demonstrated consistent improvement in edge recovery, ultimately achieving complete retrieval (HR = 1) at higher ranking positions.

### 2.5 Faithfulness and stability assessment of identified explanations

To evaluate the quality of explanations generated by DBR-X and ensure their reliability for drug repositioning, we assessed their faithfulness and stability using three established metrics: deletion, insertion, and stability [26].

The **deletion** metric assesses the importance of edges identified by DBR-X by measuring how their removal affects the model’s predictive accuracy. If the edges prioritized by the learned mask weights are truly critical, removing them should significantly disrupt the model’s ability to maintain its original predictions. To implement this, we used a sliding window approach to systematically remove edges from the query subgraph in order of decreasing mask weight. At each step, we removed 10% of the total edges (in increments of 0.2% per window) and recomputed the node representations using the trained link-prediction module on the modified subgraph. We then quantified the impact by calculating the Euclidean distance between the query disease node’s representation in the original subgraph and its representation in the modified subgraph. If the mask accurately identifies critical edges, their removal should significantly alter the query disease node’s representation, increasing the distance from its original state and indicating their essential role in the prediction.

As depicted in Figure 3d, the DBR-X illustrates an increase in distance between the disease representations of the original subgraph and the modified subgraph when the top-ranked edges, prioritized by mask weights, are removed. The rise in this distance highlights DBR-X’s superior faithfulness, reflecting its ability to isolate a concise set of highly influential edges critical for preserving predictive accuracy. While CBR-PaGE shows a similar performance, it shows a more gradual decrease, suggesting a less pronounced sensitivity to the removal of top-ranked edges. In contrast, CBR+GNNExplainer and CBR+PGExplainer exhibit flatter distance increases, indicating a less discerning prioritization of edges.

Complementing deletion, the **insertion** metric evaluates how well the model retains its predictive accuracy when only the most important edges, as ranked by mask weights, are included. This tests whether a minimal subset of high-weighted edges can sufficiently preserve the original prediction, reflecting the explanatory power of the identified connections. We began by retaining only the top 10% of edges with the highest mask weights, removing all others from the subgraph. Using the same sliding window approach (0.2% increments), we incrementally added edges back in order of decreasing importance, recomputing the node representations at each step. The Euclidean distance was calculated between the query disease node’s representation in the original complete subgraph and its representation in the modified subgraph. If the mask accurately identifies critical edges, this distance should remain small when only these edges are present, indicating that the reduced subgraph captures the essence of the original prediction.

As shown in Figure 3e, the DBR-X illustrates a sharp decrease in distance between the disease representations of the original subgraph and the modified subgraph when the top-ranked edges, prioritized by mask weights, are retained. This indicates that a small subset of high-weighted edges is sufficient to preserve the original prediction, highlighting the explanatory power of the selected connections. While CBR+PaGE shows a comparable trend as DBR-X, it exhibits a slower increase, suggesting less efficiency in capturing the most critical edges. In contrast, CBR+GNNExplainer and CBR+PGExplainer display higher initial distances and flatter decreases, indicating a less discerning prioritization of edges essential for maintaining the original prediction.

Next, **stability** measures the robustness of explanations against structural perturbations, a crucial property for ensuring reliability in real-world applications where biological networks may contain noise or incomplete data. We introduced random edge additions to each query subgraph, reran the explanation algorithms, and computed the Pearson correlation between the original and perturbed mask weights. A high correlation indicates that the explanation remains consistent despite modifications. As depicted in Figure 3f, DBR-X maintained a consistently high correlation 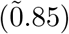 across varying levels of perturbation, outperforming all baselines. While CBR+PaGE also showed reasonable stability 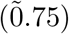, DBR-X’s higher and more consistent correlation values demonstrate greater robustness of its explanations under random modifications to the query subgraph.

### 2.6 Validating DBR-X rationales against medical evidence

Among the approximately 7,000 to 10,000 known rare diseases, only 4-6% have FDA-approved treatments, leaving the vast majority without viable therapeutic options [27]. To evaluate DBR-X’s potential for addressing this significant therapeutic gap, we examined its predictions and explanatory pathways for three rare diseases that exemplify distinct therapeutic areas. We first used DBR-X’s link prediction module to identify potential repositioning candidates from the top-10 ranked predictions. We then employed the path identification module to derive the most important biological mechanism connecting each drug to its predicted disease target. Lastly, we examined their multi-hop rationales relative to existing biomedical knowledge.

First, we examined DBR-X prediction for Duchenne muscular dystrophy (DMD), a genetic disease characterized by progressive muscle weakness due to mutations in the dystrophin gene [28]. DBR-X predicted bitolterol as a potential therapeutic agent, specifically targeting the *β*2-adrenergic receptor (*ADRB2*) (Figure 4a). Bitolterol is traditionally prescribed as a bronchodilator to treat breathing difficulties, acting as an agonist of *β*2-adrenergic receptors. Extensive scientific literature demonstrates that *β*2-adrenergic receptor agonists can stimulate skeletal muscle hypertrophy by maintaining the rate of muscle protein synthesis and/or degradation [29–31]. The activity of these receptors is tightly controlled through clathrin-mediated endocytosis, which regulates their availability on the cell surface [26]. This predicted mechanism suggests that bitolterol could help combat the muscle-wasting characteristic of DMD by maintaining muscle mass, potentially slowing disease progression.

**Fig. 4.**
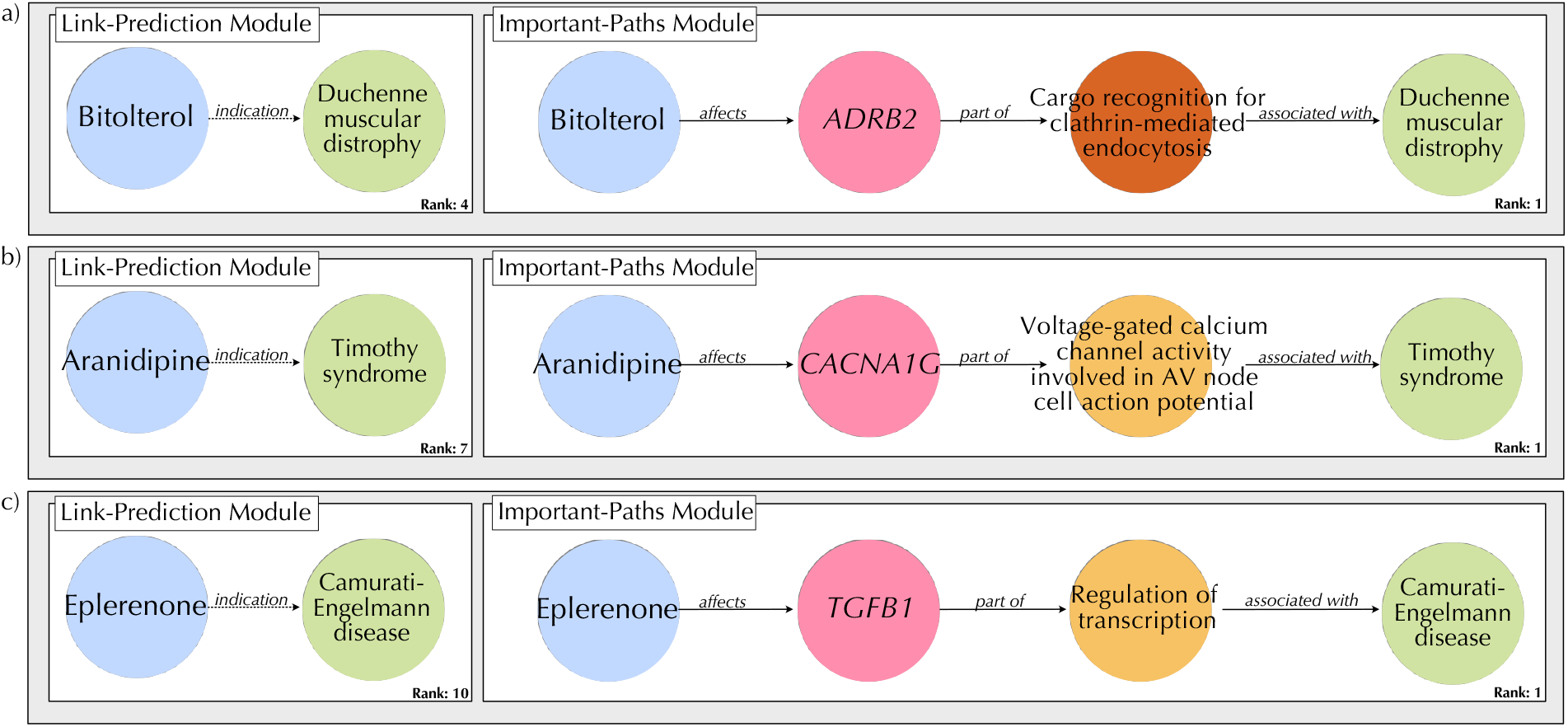
Validation of drug-repositioning candidates identified by DBR-X for three rare diseases based on medical evidence. (A) Bitolterol is predicted as a therapeutic candidate for Duchenne muscular dystrophy (rank 4), operating through *ADRB2* -mediated regulation of clathrin-mediated endocytosis. (B) Aranidipine is identified as a potential treatment for Timothy syndrome (rank 7), acting via *CACNA1G*’s role in voltage-gated calcium channel activity and AV node cell action potential. (C) Eplerenone is suggested as a therapeutic option for Camurati-Engelmann disease (rank 10), functioning through *TGFB1* -mediated transcriptional regulation.

We next investigated DBR-X predictions for Timothy syndrome, a rare genetic disorder affecting the heart. This condition is caused by mutations in the *CACNA1C* gene, which encodes the CaV1.2 L-type voltage-gated calcium channel, and manifests as atrioventricular block of AV cells, prolonged QT interval, and syndactyly [32]. DBR-X identified aranidipine, an antihypertensive calcium channel blocker, as a potential therapeutic option through its interaction with the T-type calcium channel *CACNA1G* (Figure 4b). While aranidipine typically functions by blocking both L-type and T-type calcium channels in vascular smooth muscle cells to lower blood pressure, the *CACNA1C* mutations in Timothy syndrome may reduce its effectiveness at L-type channels. Nevertheless, the drug’s ability to modulate T-type channels like *CACNA1G* suggests a promising alternative mechanism for treating the cardiac manifestations of Timothy syndrome, potentially helping to prevent atrioventricular block and normalize QT intervals.

In the final example, we looked at Camurati-Engelmann disease (CED), a rare genetic disorder affecting bone growth and development due to mutations affecting transforming growth factor-*β*1 (*TGFB1*). Under normal conditions, *TGFB1* maintains bone homeostasis by coordinating the activities of osteoblasts and osteoclasts. However, when mutations disrupt *TGFB1* function, patients experience progressive bone thickening, leading to chronic pain and increased fracture risk [33] . DBR-X identified eplerenone as a promising therapeutic candidate (Figure 4c). Eplerenone, currently used as an aldosterone receptor antagonist, has been shown to prevent atrial fibrosis by modulating the *TGFB1* signaling pathway [34]. This prediction aligns with current medical understanding, as studies in mouse models have demonstrated that targeted regulation of TGF-*β*1 signaling can help maintain healthy bone formation patterns [35].

## 3 Discussion

Understanding drug mechanisms of action remains a critical challenge in drug repositioning. Biomedical knowledge graphs have emerged as powerful tools for integrating diverse biological relationships into structured networks, from protein-protein interactions to drug-target bindings and disease associations. While GNNs have shown promise in analyzing these knowledge graphs to predict novel connections, their “black box” nature makes mechanistic understanding difficult, limiting their value in clinical applications. Existing explainable GNN approaches attempt to address this interpretability gap, but often produce explanations that fail to align with biological reality. DBR-X addresses these limitations by combining case-based subgraph analysis with heterogeneous path-enforcing mask learning, leveraging the principle that similar drugs likely act through similar biological pathways to identify both novel therapeutic targets and their underlying mechanisms.

The improved performance of DBR-X over other GNN-based link prediction frameworks demonstrates that focusing on query-specific subgraphs derived from similar cases, rather than considering the entire graph during message-passing, leads to more accurate predictions. While all models utilize graph neural architectures for learning representations, they differ fundamentally in their message-passing strategy. DBR-X employs a R-GCN strategy for encoding local structures within query-relevant subgraphs identified through case-based reasoning, whereas baseline methods perform message passing across the entire knowledge graph. By focusing only on the most relevant biological pathways for each query, DBR-X effectively filters out noise and captures more complex biological relationships. This context-specific strategy offers several critical advantages for drug repositioning. First, by restricting analysis to subgraphs from similar drug cases, DBR-X effectively filters out noise from the vast network of biological interactions, focusing only on pathways relevant to the therapeutic context of interest. Second, the case-based framework leverages successful drug applications as templates, enabling the model to identify similar therapeutic patterns that may be applicable to new diseases. Third, analyzing the clustering of diseases in the representation space helps identify previously unrecognized therapeutic opportunities where drugs might be repurposed based on shared biological mechanisms.

DBR-X provides a ranked list of multi-hop interpretable biological explanations through its path-identification module, offering detailed mechanistic rationales for predicted drug-disease associations. The core challenge of identifying these explanations lies in appropriately weighing the contributions of different biological entities within the network. To address this, DBR-X employs a node degree scoring mechanism that systematically evaluates each node’s importance based on its network connectivity, recognizing that biological networks contain both highly connected hub nodes and more specialized entities that serve distinct functional roles. As shown in the result section, optimal performance occurs with moderate degree values, avoiding the extremes of both highly connected and isolated nodes. This finding aligns with fundamental biological principles: highly connected nodes, often represent general cellular processes that may not be specific enough for drug targeting. On the other hand, sparsely connected nodes might represent noise in the network, or highly specialized interactions that occur under specific conditions.

DBR-X has several important limitations that should be acknowledged and addressed in future work. First, the pre-computed similarity matrix used for initial drug case retrieval relies on a simple inner-product of connectivity patterns, which may not capture the full complexity of drug relationships. Second, the current reliance on existing mechanistic knowledge in DrugMechDB for validation may constrain our ability to identify novel mechanistic pathways. We envision a comprehensive set of solutions to address these limitations. First, integration of diverse drug similarity metrics combining molecular structure fingerprints and pharmacological properties. Second, development of more sophisticated validation approaches that incorporate emerging experimental evidence and real-world clinical outcomes data to supplement existing mechanistic databases. We also envision extension of the framework to incorporate tissue-specific network, which would allow DBR-X to capture context-dependent drug effects and potentially lead to more precise predictions for specific disease contexts and patient populations.

In conclusion, DBR-X offers not just predictions but also clear mechanistic hypotheses that can be experimentally validated. This transparency is crucial for building trust in AI-assisted drug discovery and accelerating the translation of computational predictions into clinical applications.

## 4 Methods

### Notation

A Knowledge Graph (KG) is a structured representation of information, and is formally defined as a directed labeled multi-graph *G* = (*V, E*), where *V* is a set of nodes and *E* is a set of edges. Facts in a KG are represented as triplets (*v*_1_, *e*_1_, *v*_2_), with *v*_1_, *v*_2_ ∈ *V* and *e*_1_ ∈ *E*, where each node *v* ∈ *V* has a type *ϕ*(*v*) ∈ *A* (with *A* being the set of possible node types) and each edge *e* ∈ *E* has a type *τ* (*e*) ∈ *R* (with *R* being the set of possible edge types). The subset of edges with type *r* ∈ *R* is denoted *E*_*r*_ = {*e* ∈ *E* | *τ* (*e*) = *r*}, which groups relationships for analysis. A path *p* in the KG is a sequence of nodes connected by edges, formally written as *p* = (*v*_1_, *e*_1_, *v*_2_, …, *e*_*n*_, *v*_*n*+1_), while a meta-path *m* abstracts this into a sequence of node and edge types, defined as *m* = (*ϕ*(*v*_1_), *τ* (*e*_1_), *ϕ*(*v*_2_), …, *τ* (*e*_*n*_), *ϕ*(*v*_*n*+1_)), capturing the structural pattern of the path.

### Problem definition

In this work, we address a *post-hoc* and instance-level GNN explanation problem. DBR-X takes a trained GNN model (Ψ) and doesn’t change the model architecture or parameters to generate explanations for the predictions of each instance (*v*_1_, *e*_1_, *v*_2_). For this, we first train a GNN model to solve the task of predicting links between known drug and disease entities (link-prediction module). Then, DBR-X answers the question of *why* a particular drug entity is predicted by Ψ to be linked with the relation “*indication*” to a particular disease (important-paths identification module).

### Method overview

DBR-X has two modules. (**i**) Link prediction: given a drug query input *q*, DBR-X retrieves *k*-nearest neighbor drug cases (*k*-NN_*q*_). For each retrieved case *c*, a collection of reasoning chains *p* leading to its corresponding disease answers are collected. Next, paths retrieved are reused on the input query to form a subgraph (*G*_*i*_), where a GNN encodes the underlying subgraph structure into a node representation. If the CBR hypothesis holds true, the answer nodes representation in query subgraph *G*_*i*_ and *k*-NN_*q*_ drug’s subgraph (*G*_*j*_) will be similar, as the local structure around them share similarities. Therefore, DBR-X identifies the answer node of a given drug query that has the most similar representation to the answer nodes in the subgraph of *k*-NN_*q*_. (**ii**) Important paths identification: For a given *G*_*i*_ that connects a drug and a disease entity, DBR-X learns a heterogeneous path-enforcing mask that identify important paths to explain the connection between *v*_1_ and *v*_2_.

### Proposed method

This section details the two DBR-X modules. Step 1: The link prediction module that follows the three main steps of the CBR framework; retrieving similar cases, reusing solutions, and revising predictions.Step 2: The heterogeneous path-enforcing mask learning module to identify important paths. A pseudo-code of DBR-X link-prediction module and important-paths module is shown in Algorithm 1 and Algorithm 2, respectively.

**Step 1:** Case-Based Reasoning for drug repositioning.

- **Retrieve similar drug cases** . Given a drug query, DBR-X first retrieves *k*-NN_*q*_ similar drug cases *c* using a pre-computed similarity matrix (*S* ∈ ℝ^*V* ×*V*^) that stores the similarity score between all pairs of drug nodes on the biomedical KG. To model this, each drug entity is parameterized in an *m*-hot vector (*v* ∈ ℝ^|*E*|^), with dimensions equal to the number of relation types on the KG. An entry in the vector is set to 1 if a drug entity has at least one edge with that relation type, otherwise is set to 0. The similarity score between two drug queries is given by the cosine similarity between their normalized vector representations. Naturally, two drug nodes that capture the same relation types should have a high similarity score. For each retrieved drug case, DBR-X gathers the paths in the graph that connect the drug entity to the corresponding disease it treats. Note that since the number of collected paths between two nodes can grow exponentially, we only considered 1,000 randomly sampled paths of length up to three around each drug.
- **Reuse drug solutions** Next, the gathered paths *p* of retrieved similar drug cases are reused for the query of interest. Beginning with the drug query node, the meta-path *m*, defined as the sequence of edge types connecting node types, is applied. An example is depicted in Figure 1a, where astemizole is retrieved as one of the *k*-NN_*q*_ similar drugs to the Pranlukast query. The recovered path *p*= Astemizole, *affects*, HRH4, *part of*, Histamine receptor, *associated*, Rhinitis is reused by deriving its meta-path *m* = Drug, *affects*, Gene, *part of*, Pathway, *associated*, Disease, which captures the sequence of relation types. After iterating over all similar retrieved cases, the collected paths form a subgraph for the drug query *G*_*i*_.
- **Revise disease predictions** Once the subgraphs of the drug query and the corresponding similar drugs are defined, DBR-X reasons across them. These subgraphs are subsets of the Knowledge Graph *G* = (*V, E*), capturing relevant nodes and edges for analysis. For this, the local subgraph structure of both the drug query and its *k*-NN_*q*_ similar drugs is encoded with a GNN. Considering that biomedical KGs are heterogeneous graphs with labeled edges, where each edge *e* ∈ *E* has a type *τ* (*e*) ∈ *R*, we employed the multi-relational R-GCN model. We followed the general message-passing neural network scheme that iteratively updates the representation of each node by aggregating the representations of its immediate outgoing neighbors. In particular, the general GNN message-passing process at the *l*^th^ layer is given by:

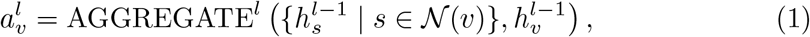

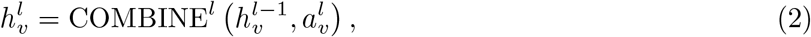

where 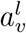 is the aggregated message from the neighbors, 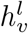 denotes the representation of node *v* in the *l*^th^ layer, and 𝒩(*v*) denotes the set of immediate outgoing neighbors of node *v*. For the multi-relational R-GCN model, which accounts for edge types, these steps are specialized as:

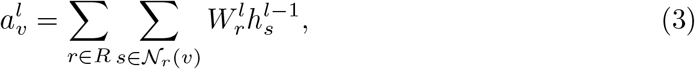

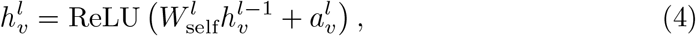

where *R* is the set of edge types captured in the KG, 𝒩_*r*_(*v*) denotes the immediate outgoing neighbors of node *v* under edge type *r*, derived from edges in *E*_*r*_ = {*e* ∈ *E* | *τ* (*e*) = *r*}, 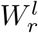 is the transformation matrix used to propagate the message in the *l*^th^ layer for edge type *r*, and 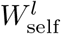 is the self-loop transformation matrix that updates the node’s own representation.

During training, the disease answer node *d*_*i*_ of the query subgraph *G*_*i*_ and the disease answer node *d*_*i*_ of the *k*-NN_*q*_ subgraphs *G*_*j*_ are trained to be more similar to each other in comparison to incorrect disease answer nodes. This similarity is calculated as the similarity between the normalized disease answer representations. Considering that a retrieved similar drug case could arrive at a set of answers *D*_*j*_, we calculate the mean of the scores between *d*_*i*_ and all answer nodes in *G*_*j*_ as:

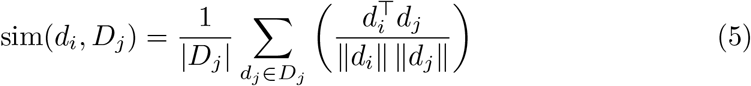

Then, the mean similarity score of all retrieved similar drug cases is aggregated for the given drug query. The loss function (ℒ_∫⟩⇕_), an adaptation of the normalized temperature-scaled cross-entropy loss from Chen et al. [36] and further developed by Das et al. [15], is defined as follows:

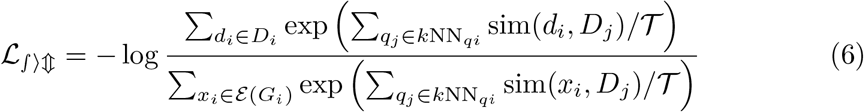

where, *D*_*i*_ represents the set of all answer nodes in *G*_*i*_, while *D*_*j*_ denotes the set of all answer nodes of retrieved similar *k*-NN_*q*_ drug cases, *x*_*i*_ ∈ *G*_*i*_ represents all nodes in the subgraph of drug query and 𝒯 denotes the temperature hyperparameter. This way, the loss score the disease answer nodes in *D*_*i*_ higher than all other nodes in *G*_*i*_ with respect to the answer nodes in *k*-NN_*q*_ drug subgraph.

Lastly, during reasoning message-passing is run across both the drug query subgraph and the subgraphs of retrieved *k*-NN_*q*_ drugs, resulting in the generation of node representations. A ranking of similarity scores is returned, where the highest score node in the query subgraph with respect to all answer nodes in the retrieved drug subgraphs is returned as a more likely repositioning answer. For this,

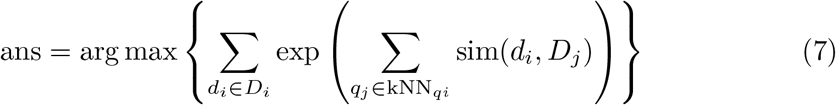

#### Algorithm 1 Pseudocode for the Link-Prediction Module of DBR-X

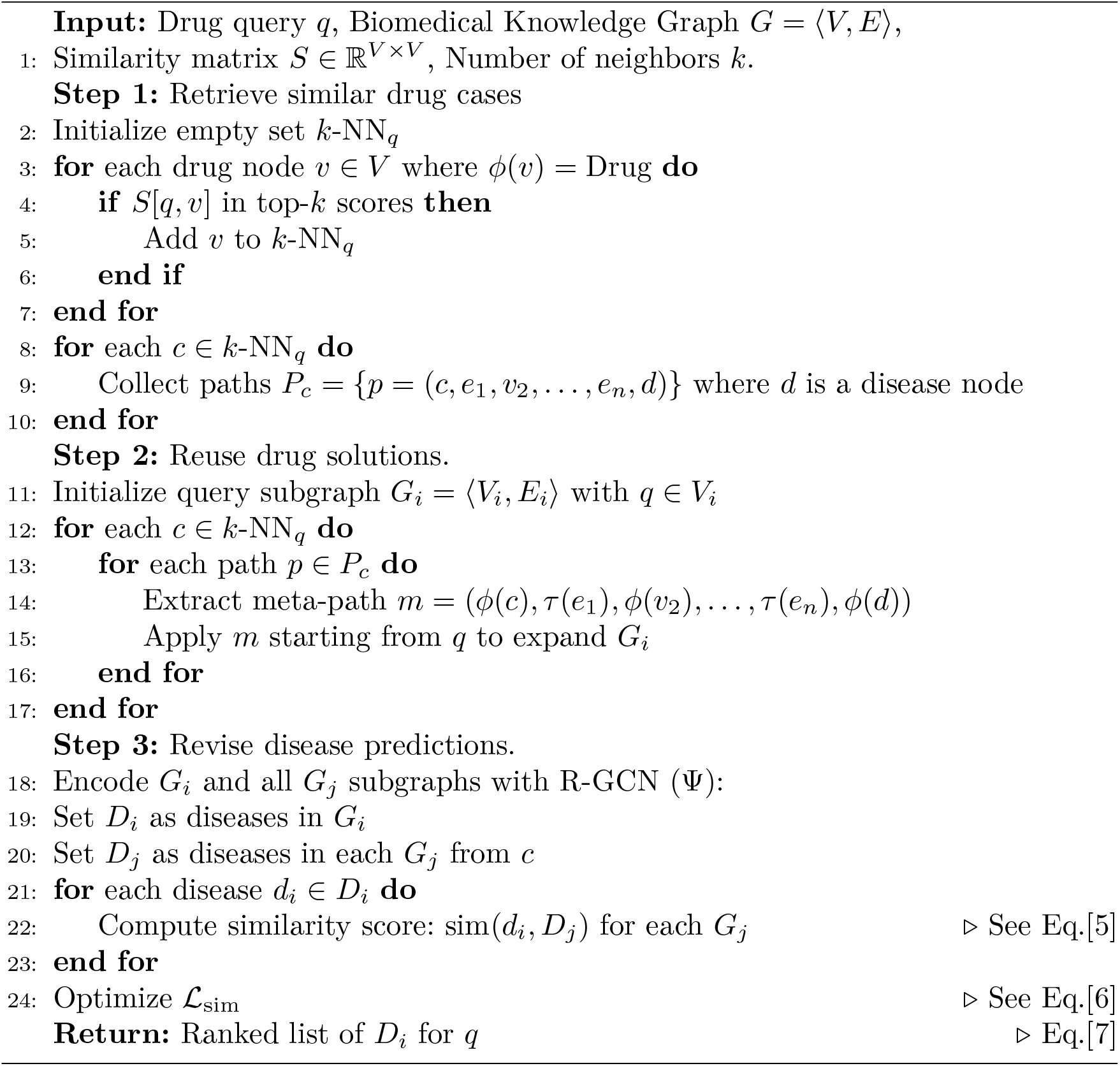

**Step 2:** Heterogeneous path-enforcing mask learning.

The second module of DBR-X learns a heterogeneous path-enforcing mask *M* to identify critical edges within the query subgraph *G*_*i*_ = ⟨*V*_*i*_, *E*_*i*_⟩ that explain the predicted drug-disease association (*q, d*). Specifically, the heterogeneous mask is defined as 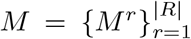, where 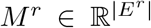 corresponds to the mask weights for edges of type *r*, and *R* is the set of edge types in the knowledge graph. To achieve this, we define the explanation graph *G*_*e*_ = ⟨*V*_*e*_, *E*_*e*_, *M* ⟩ as a weighted subgraph of *G*_*i*_, where *V*_*e*_ ⊆ *V*_*i*_ includes the drug node *q*, the disease node *d*, and all nodes along paths connecting them, and *E*_*e*_ ⊆ *E*_*i*_ consists of edges assigned weights by the mask 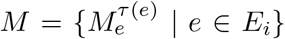. Each edge *e* ∈ *E*_e_ has a weight 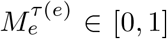, reflecting its importance to the prediction, with higher weights indicating greater influence on the GNN’s message-passing process. Initially, *G*_*e*_ is set to *G*_*i*_ with a random mask *M* ^(0)^, where each *M* ^*r*(0)^ is initialized randomly for all edge types *r* ∈ *R*, and during training, *M* is iteratively updated to retain edges that both preserve the predictive accuracy of the link-prediction module and form biologically meaningful paths. The goal is to refine *G*_*e*_ such that it captures the most explanatory subgraph connecting *q* to *d*.

Here, mask learning for link prediction explanation is done from two perspectives: important edges should be influential for the link-prediction module, and form meaningful paths. For this, we introduce the loss Loss_explanation_ that allows us to achieve these two measurements:

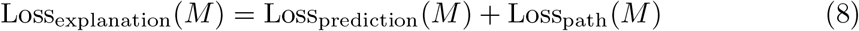

First, *L*_*prediction*_ is the loss term for *M* to learn to select influential edges for model prediction. The idea is to perform a perturbation-based explanation, where parts of the input are considered important if perturbing them significantly changes the model’s prediction. If removing an edge alters the original similarity between the answer node *d*_*i*_ of the query subgraph *G*_*i*_ and the disease answer node *d*_*j*_ of the *G*_*j*_ subgraphs, then the edges is a critical counterfactual edge that should be part of the explanation. This idea can be formalized as minimizing the similarity difference between the original disease answer node representation and the disease answer representation of the masked graph *G*_*e*_.

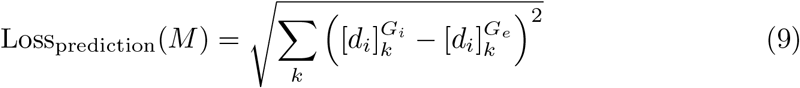

Loss_prediction_quantifies the change in the original *d*_*i*_ representation when it is limited to the explanation graph *G*_*e*_. This way, it learns to identify candidate edges by enforcing the explanation graph to keep the original representation.

Next, Loss_path_ is the loss term for *M* to learn to select edges that form informative paths. The mask optimization forces weights of influential edges 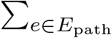 to increase, while the mask weights of non-informative edges 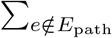 to decrease. For this, we consider the weighted average of these two terms, regulated by the hyperparameters *α* and *β*.

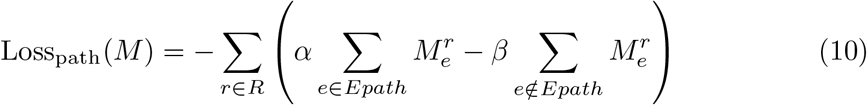

To compute Loss_path_ it is necessary to define the edges that form informative paths 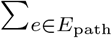 . For this, we define the importance of a path with the score function Score(*p*). Each edge score is defined by the probability of including *e* in the explanation 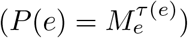 and a node degree score *D* .

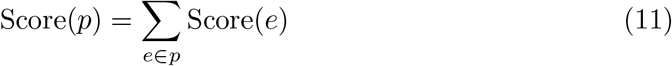

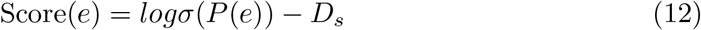

We define the node degree score as *D*_*s*_ = *log*(*κ* + *d − γ*), that measures how far a target node’s degree (*d*) is from a specified degree score (*γ*). The constant *κ* is included to ensure that the argument of the logarithm is always positive, avoiding undefined or negative values. By applying the logarithm function, we achieve a normalized scale that mitigates the impact of large deviations and effectively handles a wide range of degree values. This scoring method provides a robust score of how closely a node’s degree aligns with a defined *γ* score. A Score(*p*) will be high if the edges on it have high probabilities and these edges are linked to nodes with a high *D*_*s*_, which would correspond to those that are closer to the defined *γ*. We employ Dijkstra’s shortest-path algorithm to identify paths with the highest Score(*p*), selecting their edges as *E*_path_. After mask learning converges, we apply Dijkstra’s algorithm once more using the final mask *M* to generate and rank the top explanatory paths.

#### Algorithm 2 Pseudocode for the Important-Paths Module of DBR-X

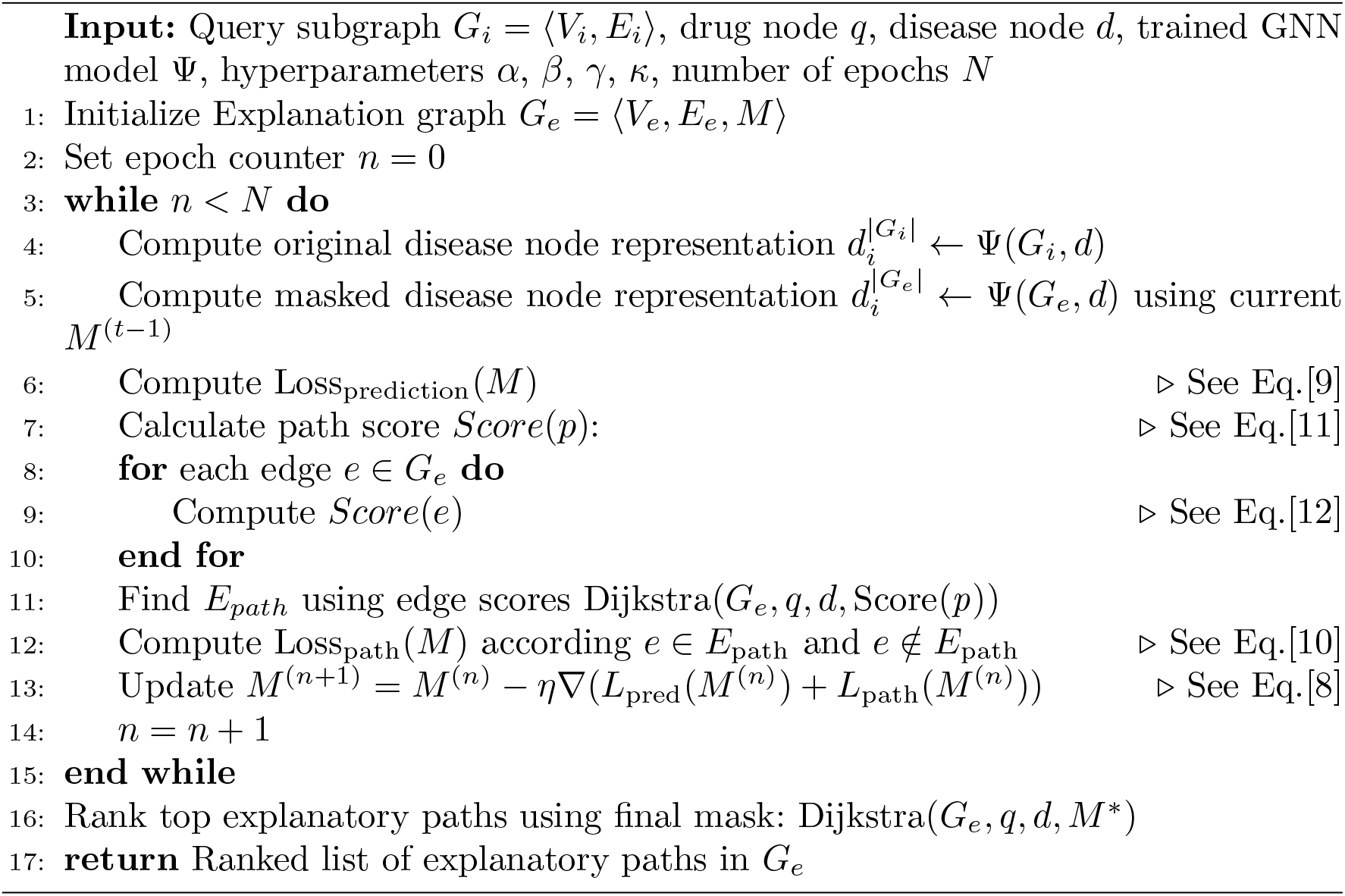

### Dataset

We conducted experiments on Mechanistic Repositioning Network with Indications (MIND), a biomedical knowledge graph that integrates two biomedical resources: Mechanistic Repositioning Network (MechRepoNet) [4] and DrugCentral [37]. Briefly, MechRepoNet is a comprehensive biomedical knowledge graph that was constructed by integrating 18 different data sources, consisting of of 9,652,116 edges, 250,035 nodes, 9 node types and 22 relations. DrugCentral, is a publicly available online resource that incorporates information from indications that have received approval from regulatory agencies. Here, knowledge graph completion prediction performance was evaluated on a subset of DrugCentral indications, for this we divided MIND into subsets: train (80%, 2087 indications), and test (20%, 390 indications).

### Hyperparameters

We conducted hyperparameter tuning using Optuna software. For the link-prediction module, the final configuration includes a GCN layer with 64-dimensional input features per node and a dropout rate of 0.7966 to randomly skip neighbor connections during aggregation, producing 128-dimensional output features. We set the number of neighbors to 5 for training and 10 for evaluation. The model was optimized using the Adam Optimizer with a learning rate of 0.1, a temperature of 0.1053, and a sampling loss weight of 0.3391.

### Implementation details

The DBR-X model is built using the DGL [38] and PyTorch [39] deep learning frameworks in Python. For data processing and computation, we utilize Pandas [40] and NumPy [41]. Evaluation metrics are handled with scikit-learn [42], while visualization is performed using seaborn [43], matplotlib [44], and UMAP [45]. Training progress is monitored using Weights and Biases [46]. The model is trained on a server equipped with a single NVIDIA Tesla V100 GPU.

## Data availability

The MIND knowledge graph can be found at https://doi.org/10.5281/zenodo.8117748. The DrugMechDB relevant files are hosted at https://doi.org/10.5281/zenodo.8139357.

## Code availability

The python code to reproduce results, documentation and usage examples is available on GitHub at https://github.com/SuLab/DBR-X.

## Acknowledgments

The authors thank Dr. Chunlei Wu and Dr. Rajarshi Das for their feedback.

## Notes

https://github.com/SuLab/DBR-X

### Competing Interest Statement

The authors have declared no competing interest.

